# Investigating the importance of anatomical homology for cross-species phenotype comparisons using semantic similarity

**DOI:** 10.1101/028449

**Authors:** Prashanti Manda, Christopher J. Mungall, James P. Balhoff, Hilmar Lapp, Todd J. Vision

## Abstract

There is growing use of ontologies for the measurement of cross-species phenotype similarity. Such similarity measurements contribute to diverse applications, such as identifying genetic models for human diseases, transferring knowledge among model organisms, and studying the genetic basis of evolutionary innovations. Two organismal features, whether genes, anatomical parts, or any other inherited feature, are considered to be homologous when they are evolutionarily derived from a single feature in a common ancestor. A classic example is the homology between the paired fins of fishes and vertebrate limbs. Anatomical ontologies that model the structural relations among parts may fail to include some known anatomical homologies unless they are deliberately added as separate axioms. The consequences of neglecting known homologies for applications that rely on such ontologies has not been well studied. Here, we examine how semantic similarity is affected when external homology knowledge is included. We measure phenotypic similarity between orthologous and non-orthologous gene pairs between humans and either mouse or zebrafish, and compare the inclusion of real with faux homology axioms. Semantic similarity was preferentially increased for orthologs when using real homology axioms, but only in the more divergent of the two species comparisons (human to zebrafish, not human to mouse), and the relative increase was less than 1% to non-orthologs. By contrast, inclusion of both real and faux random homology axioms preferentially increased similarities between genes that were initially more dissimilar in the other comparisons. Biologically meaningful increases in semantic similarity were seen for a select subset of gene pairs. Overall, the effect of including homology axioms on cross-species semantic similarity was modest at the levels of divergence examined here, but our results hint that it may be greater for more distant species comparisons.

## 1. Introduction

### 1.1. Cross-species phenotype matching

Organisms exhibit similarities with each other in their genetic content, anatomical structures, and other biological features due in large part to common evolutionary descent. This similarity is what allows non-human organisms to serve as models for human diseases and for biological knowledge to be transferred from model organisms to related species. In the area of biomedical informatics, an important recent application is the use of cross-species phenotype matching algorithms to generate candidate gene lists for rare and undiagnosed diseases.^1,2^ Given a phenotypic profile for a human disease (e.g. a list of terms from The Human Phenotype Ontology,^3^ cross-species profile matching tools generate a ranked list of candidate genes based on matches to the phenotypic profiles of orthologous genes in mutant models. This process can be automated by using phenotype ontologies and semantic similarity methods that quantify the degree of similarity.^4,5^ A number of methods make use of the Uberon anatomy ontology to connect phenotype terms across species.^6,7^ For example, the human phenotype “Abnormality of the upper limb” (HP_0002817) is connected to the mouse phenotype “abnormal forelimb morphology” (MP_0000550) via the Uberon class “forelimb” (UBERON_0002102).

### 1.2. Homology

Two organismal features, whether genes, anatomical parts, or another inherited feature, are considered to be homologous when they are evolutionarily derived from a single feature in a common ancestor. Orthologous genes are a particular class of homologous features, ones that are found in two different organismal lineages and that split evolutionarily into two genetic lineages during a speciation event. It is a foundational premise for much of comparative genomics that orthologous genes retain comparable functions even in distantly related organisms.^8^ For example, in chick, *Tbx5* and *Tbx4* genes control early development of wing and hindlimb buds respectively, and the orthologs of these genes in zebrafish control development of *anatomically homologous* structures, the pectoral and pelvic fins^9^ (Figure 1). Thus, it appears that these two gene lineages were distinct in the common ancestor of fish and birds and were deployed similarly in the development of the ancestral fore and hind appendages.

**Fig. 1.**
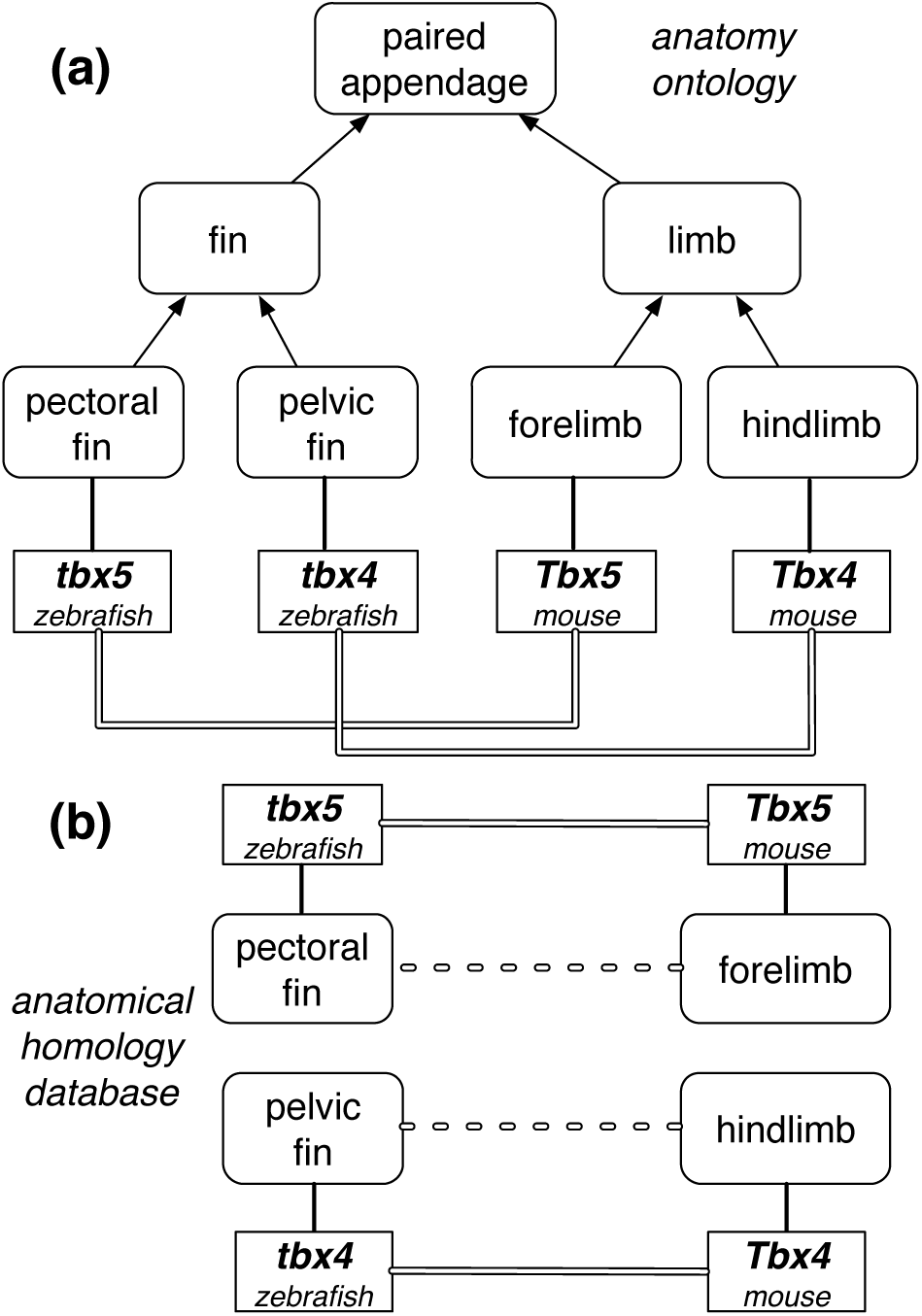
The role of an orthologous gene pair in the development of different appendages. (a) genes (square boxes) are expressed in anatomical structures (rounded boxes), which are organized hierarchically in a subclass hierarchy (arrows). The two ortholog pairs are anatomically similar only at the level of “paired appendage”. (b) Adding anatomical homology (dotted lines) increases anatomical similarity between orthologous pairs.

Recognizing similar phenotypes grows increasingly challenging as the evolutionary distance increases between species and anatomical features diverge in structure. Comparative anatomists have given a great deal of attention to identifying homologous anatomical structures among distantly related species.^10^ Uberon does not contain explicit homology relationships,^11^ such as between a hindlimb and a pelvic fin, or between the mammalian adrenal gland and the zebrafish interrenal gland. Instead, these classes are grouped according to similar structure, function or cellular composition. For example, both *forelimb* and *hindlimb* are grouped under the more general class *limb* based on their shared morphology, and limbs and fins are grouped under the general *paired appendage* (Figure 1). The situation is complicated by the fact that homology assumptions do necessarily leak into the construction of the ontology. The fact that the forelimb and hindlimb are similar morphologically is no accident if it is accepted that these are anatomical serial homologs. In fact, Uberon includes a grouping class *paired limb/fin* (UBERON_0004708) based on homology. Despite the above, homologous structures may sometimes be placed relatively distant to each other within Uberon when structural similarities are not as apparent (e.g. as is the case for certain bones in the jaw of fish that are homologous to the inner ear bones of mammals). Phenotypes affecting anatomical features that are homologous, but distantly placed within the ontology, will appear artificially dissimilar to one another.

We wish to quantify the extent to which the accuracy of cross-species phenotype matching is increased by including assertions of homology, such as those compiled by Bgee,^12^ into Uberon. We do this by assessing how measures of semantic similarity are affected for orthologous relative to non-orthologous gene pairs. The underlying assumption is that orthologous genes are more often expressed in, and thus contribute to phenotypes in, homologous anatomical structures than non-orthologous genes, independently of how close the anatomical structures are within Uberon. If this assumption is correct, and homologies do indeed contribute to accuracy, we would expect to see a relatively greater increase in semantic similarity for orthologous genes relative to non-orthologous genes when non-trivial homologies are added to Uberon.

## 2. Methods

### 2.1. Phenotype Annotation Data

Orthologous gene pairs in zebrafish, mouse, and human were obtained from PANTHER (06/19/2014 release, v9.0).^13,14^ Phenotype annotations for these genes were obtained from the Monarch Initiative v1.0 release (https://github.com/monarch-initiative/monarch-owlsim-data), which aggregates data from the Human Phenotype Ontology (HPO), Mouse Genome Informatics (MGI), and Zebrafish Information Network (ZFIN) (see http://monarchinitiative.org for details).

Gene pairs in which one or both genes lacked phenotype annotations were not included in the final analysis. We removed 503 mouse genes whose annotations indicate a lack of phenotypic assay (MP_0003012: no phenotypic analysis) and/or the absence of an abnormal phenotype (MP_0002169: no abnormal phenotype detected). Zebrafish and human genes for which no phenotypes are presently annotated were prefiltered by the source model organism databases (ZFIN, HPO). Anatomical homology axioms for Uberon classes were obtained from the GitHub repository of Bgee v0.2 (https://github.com/BgeeDB/anatomical-similarity-annotations/blob/master/release/raw_similarity_annotations.tsv).^15^ Non-orthologous gene pairs across zebrafish, mouse, and human were randomly sampled with a uniform distribution from the set of gene pairs not asserted to be orthologous by PANTHER.

There are a variety of different semantic similarity measures used in the bioinformatics literature.^16^ Here, we present results for a commonly used measure, *Sim_IC_*, which is based on the concept of *Information Content* (IC), or the specificity of the match between two annotations relative to a chosen annotation corpus.^17^ We also examined another commonly used measure, Jaccard similarity (*Sim_J_*) which measures the ontological graph overlap between two annotations.^18^ They differ in that *Sim_IC_* takes into account the distribution of annotations among ontology terms while *Sim_J_* considers ontology structure independent of annotation density. These metrics were compared because of their prior use as measures of phenotypic similarity between orthologous genes.^2^

We also have a choice in how to summarize the set of pairwise semantic similarities between two genes, both of which typically have multiple annotations. We refer to the union of the individual phenotype annotations for all alleles of one gene as a *phenotype profile.* Here, we evaluated two summary statistics for semantic similarity between two phenotype profiles, as detailed below, which we call Best Pairs and All Pairs. We only report full results for one combination of statistics, *Sim_IC_* with Best Pairs, based on a test for which combination best discriminated between orthologs and non-orthologs (see Results).

The *IC* of ontology graph node *N* in an annotation corpus with *Z* genes is defined as the negative logarithm of the probability of a gene being annotated to *N*.

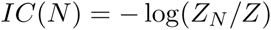

where *Z_N_* is the number of genes annotated to *N*. The *IC* for a pair of annotations, *K* and *L*, is defined as the *IC* of their *most informative common ancestor* (MICA), which is their most specific common subsumer in the ontology. Raw IC scores range from [0, *IC_max_*], with 0 being the score of the root node of the ontology graph, and *IC_max_* = −*log*(1*/Z*) the score of a node with only one gene annotation in the dataset. To obtain an IC score with a range of [0, 1], the IC-based similarity measure, *Sim_IC_*(*K*, *L*), is normalized as follows.

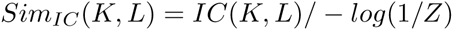

The Jaccard similarity for a pair of annotations *K, L* is defined as the ratio of the number of nodes in the intersection of their subsumers in the ontology graph over the number of nodes in the union of their subsumers.^18^

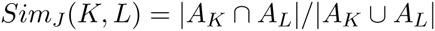

where *A_K_* and *A_L_* are the sets of subsumers of *K* and *L*, respectively.

#### 2.1.1. Similarity between phenotype profiles

To compute the Best Pairs score between two phenotype profiles *X*, *Y*, for each annotation in *X*, the best scoring match in *Y* is determined, and the median of the |*X*| values is taken. Similarly, for each annotation in *Y*, the best scoring match in *X* is determined, and the median of the |*Y*| values is taken. The Best Pairs score *S_BP_*(*X*, *Y*) is the mean of these two medians.

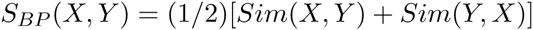

where

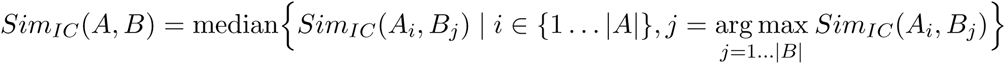

To compute the All Pairs score, one instead takes the median of of all pairwise phenotype similarities between *X* and *Y*.

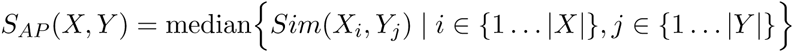

For both *S_BP_* and *S_AP_*, similarity may be measured using either *Sim_IC_* or *Sim_J_*.

### 2.2. Construction of ontologies

We constructed three ontologies for computing semantic similarity: one without homology axioms (*R*), one with valid homology axioms (*H*) and one with a random set of homology axioms (*H*′). Figure 2 illustrates the process by which these were built. Following the approach of Kohler et al.,^19^ *R*, *H* and *H*′ were seeded with the ontologies used by the gene phenotype annotations for all three species in the corpus: the mammalian phenotype,^20^ zebrafish Phenotype,^19^ and human phenotype^21^ ontologies, as well as the cross-species Uberon anatomy ontology^11^ and the phenotypic quality ontology PATO.^22^

**Fig. 2.**
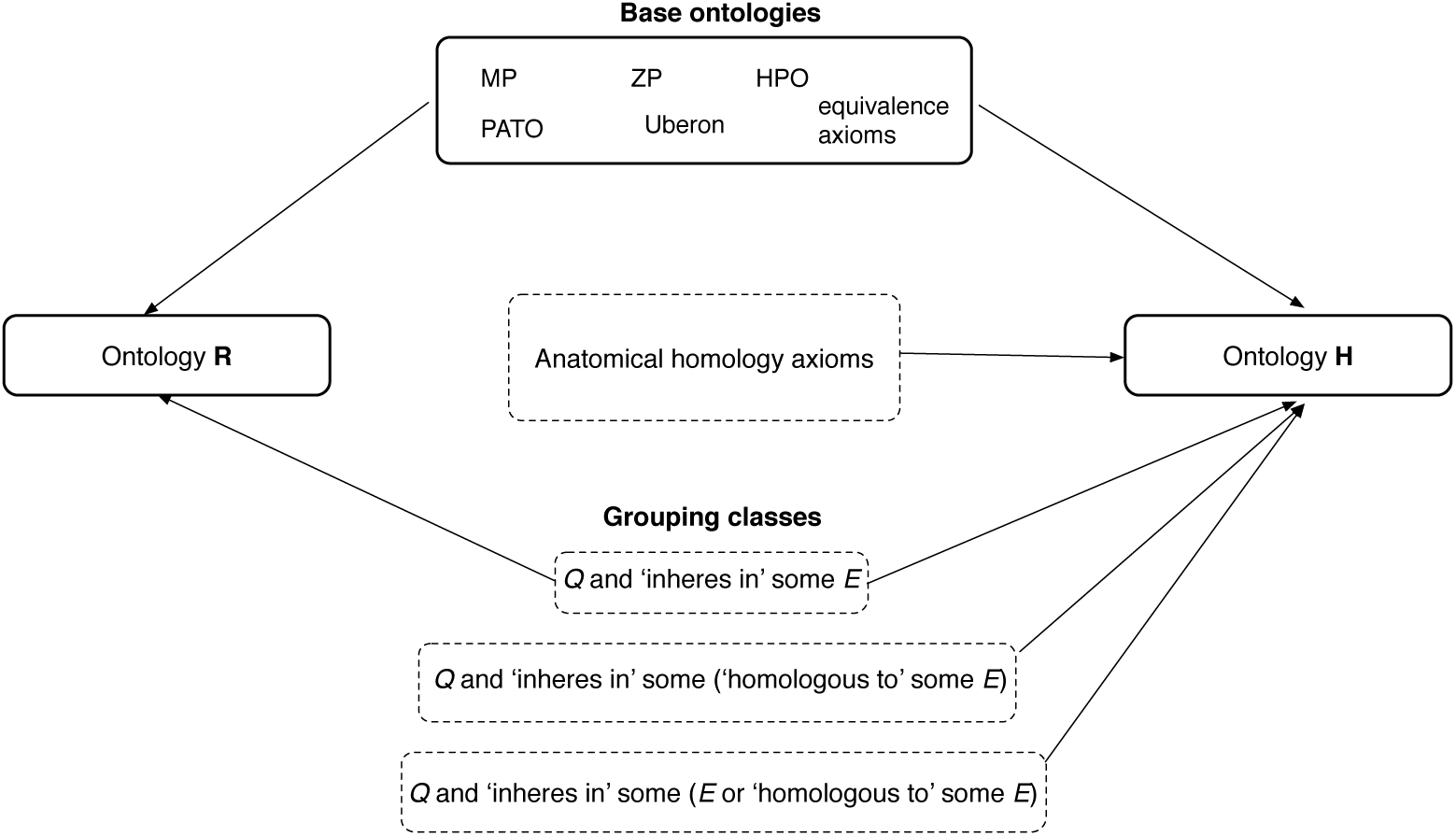
Construction of ontologies for computing semantic similarity without or with supplemented knowledge of anatomical homology. First, ontology *R* is created by adding building block phenotype ontologies and equivalence axioms. *H* (and *H*′) are created by adding anatomical homology axioms to *R*. Finally, one set of grouping classes is added to *R* and three sets of grouping classes are added to *H* (and *H*′).

There already exist a number of *homology-grouping* classes in the Uberon ontology that bundle morphologically or functionally distinct subclasses based entirely on homology. As we are seeking to determine the effect of anatomical homology on cross-species phenotype similarity of orthologous genes, we removed a number of *homology-grouping* classes from *R*, *H*, and *H*′ (Table 1).

**Table 1.** *homology-grouping* classes in Uberon excluded from *R*, *H*, and *H*′.

| Name | UBERON_ID | Name | UBERON_ID |
| --- | --- | --- | --- |
| adrenal/interrenal gland | 0006858 | paired limb/fin bud | 0004357 |
| limb/fin segment | 0010538 | paired limb/fin cartilage | 0007389 |
| paired limb/fin skeleton | 0011582 | pelvic appendage | 0004709 |
| paired limb/fin | 0004708 | pectoral appendage | 0004710 |
| paired limb/fin field | 0005732 | bone of free limb or fin | 0004375 |

Next, 1, 836 homology axioms from Bgee^15^ relating homologous anatomical structures were added to *H*. For example, “pectoral fin” is asserted as being homologous to “forelimb” and “pelvic fin” to “hindlimb”. These axioms restored the relations indicated by the excluded *homology*_*grouping* classes in Table 1. We generated a set of 1, 836 ‘random’ homology axioms by sampling anatomy terms from a permuted list of those used in the real homology axioms; these were then added to *H*′.

We then created *grouping classes* to subsume annotations based on different ontology class properties. These grouping classes are classified by an OWL reasoner into the pre-existing phenotype class hierarchy by virtue of subsumption reasoning and equivalence axioms. These axioms follow the standard Entity–Quality (EQ) template.^7,19^ In their simplest form, EQ expressions describe a phenotype in terms of a quality (*Q*) and an entity (*E*) that is the bearer of the quality. These EQ expressions are represented in OWL as “*Q* and ‘inheres in’ some *E*”.^7,23^ The following three EQ expressions were created to serve as templates for equivalence axioms of grouping classes.

- *EQ*_1_: *Q* and ‘inheres in’ some *E*
- *EQ*_2_: *Q* and ‘inheres in’ some (‘homologous to’ some *E*)
- *EQ*_3_: *Q* and ‘inheres in’ some (*E* or ‘homologous to’ some *E*)

In the above expressions, *Q* is the root of the PATO ontology and *E* can be any entity from the Uberon ontology. One class of the form *EQ*_1_ was created for each entity class (*E*) in Uberon. These classes were added to *R*, *H*, and *H*′.

The templates *EQ*_2_ and *EQ*_3_ generate classes that group annotations whose anatomical structures are related via homology. For each entity class (*E*) in Uberon, we added to ontologies *H* and *H*′ one class of form *EQ*_2_ and one of form *EQ*_3_. It is the presence of these grouping classes that control whether anatomical homology axioms are or are not used to infer common subsumers for phenotype annotations. Figure 3 illustrates the difference between ontologies for which the *EQ*_2_ and *EQ*_3_ classes are or are not included. When using *R*, which has *EQ*_1_-template grouping classes only, fin and limb phenotypes are grouped at the high level of *paired appendage.* When using *H*, the phenotypes are grouped by an *EQ*_2_-template grouping class for a more specific and thus more informative class defined by anatomical homology.

**Fig. 3.**
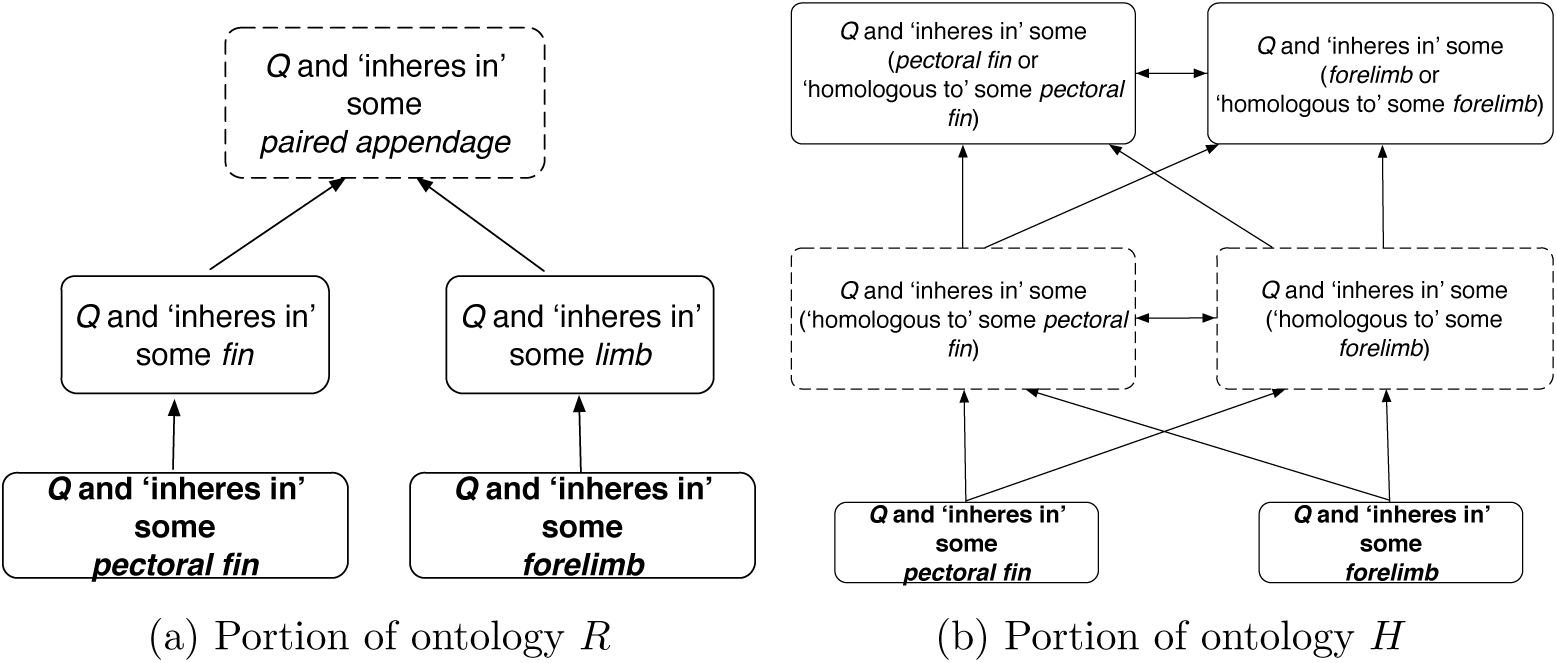
An example of subsumption hierarchies without and with anatomical homology. Least common subsumers for annotations of the pectoral fin and forelimb (dashed boxes) can be seen to differ for the *R* ontology, without homology (a) and the *H* ontology, with homology (b). Arrows denote subsumption relationships. It can be seen here that knowledge of homology enables the inference of more informative common subsumers for annotations with homologous anatomical structures.

### 2.3. Assessing the impact of homology

Our overall goal is to assess how the semantic similarity between the phenotypic profiles of two genes is affected by the addition of explicit homology statements in *H* lacking in *R*. Specifically, we hypothesized that there would be a greater increase in the similarity score for orthologous than non-orthologous genes when real homologies were included, and no differential increase when random homology assertions were included. We tested this hypothesis for two species pairs: mouse-human and zebrafish-human, with the expectation that the effect of homology on semantic similarity scores would be greater for the more distant evolutionary comparison.

To carry out this test, we measured the difference in similarity using *R* versus using either *H* or *H*′. We performed unpaired, one-sided *t*-tests for the null hypothesis that the distribution of differences was identically distributed for orthologs and non-orthologs. The alternate hypothesis is that the difference would be greater for orthologs. We performed four such tests, for both the zebrafish-human and mouse-human comparisons and for both *H* and *H*′.

## 3. Results

We obtained 1,253 orthologous gene pairs between zebrafish and mouse, 640 between zebrafish and human, and 2,034 between mouse and human, from PANTHER. Equal numbers of nonorthologous gene pairs were obtained for each species pair by sampling from permuted lists of the genes included in the orthologous pairs and requiring that the sampled pair not be included in the PANTHER orthology list. 10,055 grouping classes were added to ontology R and 30,165 to ontologies *H* and *H*′. As noted above, 1,836 anatomical homology axioms were obtained from Bgee for inclusion in *H*; they come from a wide variety of sources (Table 2). The same number of random homology axioms were added to *H*′.

**Table 2.** Sources of Bgee homology axioms classified by Evidence Code.^24^

| Evidence Code | No. axioms | Evidence Code | No. axioms |
| --- | --- | --- | --- |
| Used in automatic assertion | 709 | Morphological similarity | 66 |
| Curator inference | 361 | Traceable author statement | 55 |
| Developmental similarity | 213 | Positional similarity | 36 |
| Phylogenetic distribution | 197 | Gene expression similarity | 32 |
| Non-traceable author statement | 137 | Compositional similarity | 30 |

One of three confidence codes, “high confidence” (34.13% of axioms), “medium confidence” (61.72%), or “low confidence” (4.15%) was associated with each homology axiom by Bgee. 1,680 of the axioms assert a class to be a homolog of itself, while only 156 of the homology axioms, belonging to 12 taxonomic groups, assert homology between pairs of anatomical structures (Table 3). Thus, only a fraction of the homology axioms would be relevant for the taxonomic comparisons being made here; for example, there are only 10 non-self homology axioms that would affect comparisons between mammals.

**Table 3.** Distribution of Bgee homology axioms among taxa, excluding self-homologies.

| Taxon name | No. axioms | Taxon name | No. axioms |
| --- | --- | --- | --- |
| Vertebrata | 38 | Mammalia | 10 |
| Tetrapoda | 28 | Metazoa | 6 |
| Bilateria | 16 | Sarcopterygii | 8 |
| Chordata | 14 | Eumetazoa | 6 |
| Amniota | 10 | Gnathostomata | 8 |
| Euteleostomi | 10 | Dipnotetrapodomorpha | 2 |

We calculated phenotype semantic similarity for each orthologous and non-orthologous gene pair using the four combinations of semantic similarity measures described in the Methods above and for each of the three different ontologies, *R*, *H*, and *H*′. In order to select one semantic similarity measure for subsequent analyses, we determined which one best distinguished orthologous from non-orthologous gene pairs, reasoning that this would be an informative indicator of biological accuracy. We calculated the difference in median rank between orthologs and non-orthologs for the zebrafish-human comparison using *H*. We found that the combination of *Sim_IC_* and *S_BP_* gave the greatest discrimination between orthologous and non-orthologous gene pairs and so report results for that statistic in what follows. Full results for all four statistics, together with analysis scripts used in this study, are available from Zenodo (doi:/10.5281/zenodo.31833).

Our hypothesis that orthologs would experience a disproportionate increase in similarity when real homology axioms were used was supported by the *t*-tests in the case of the zebrafish-human comparison (Table 4). A one-tailed unpaired *t*-test found a significantly greater difference for orthologs than non-orthologs with real homology axioms but no significant difference with random axioms. However, this pattern was not seen in the mouse-human comparison, where orthologs were not significantly different than non-orthologs for *H*. In fact, the reverse trend was seen in all other comparisons; the mean similarity was preferentially increased for non-orthologs in the zebrafish-human *H*′, mouse-human *H*, and mouse-human *H*′ comparisons (Table 4). The underlying profile similarity values can be seen in Figure 4.

**Fig. 4.**
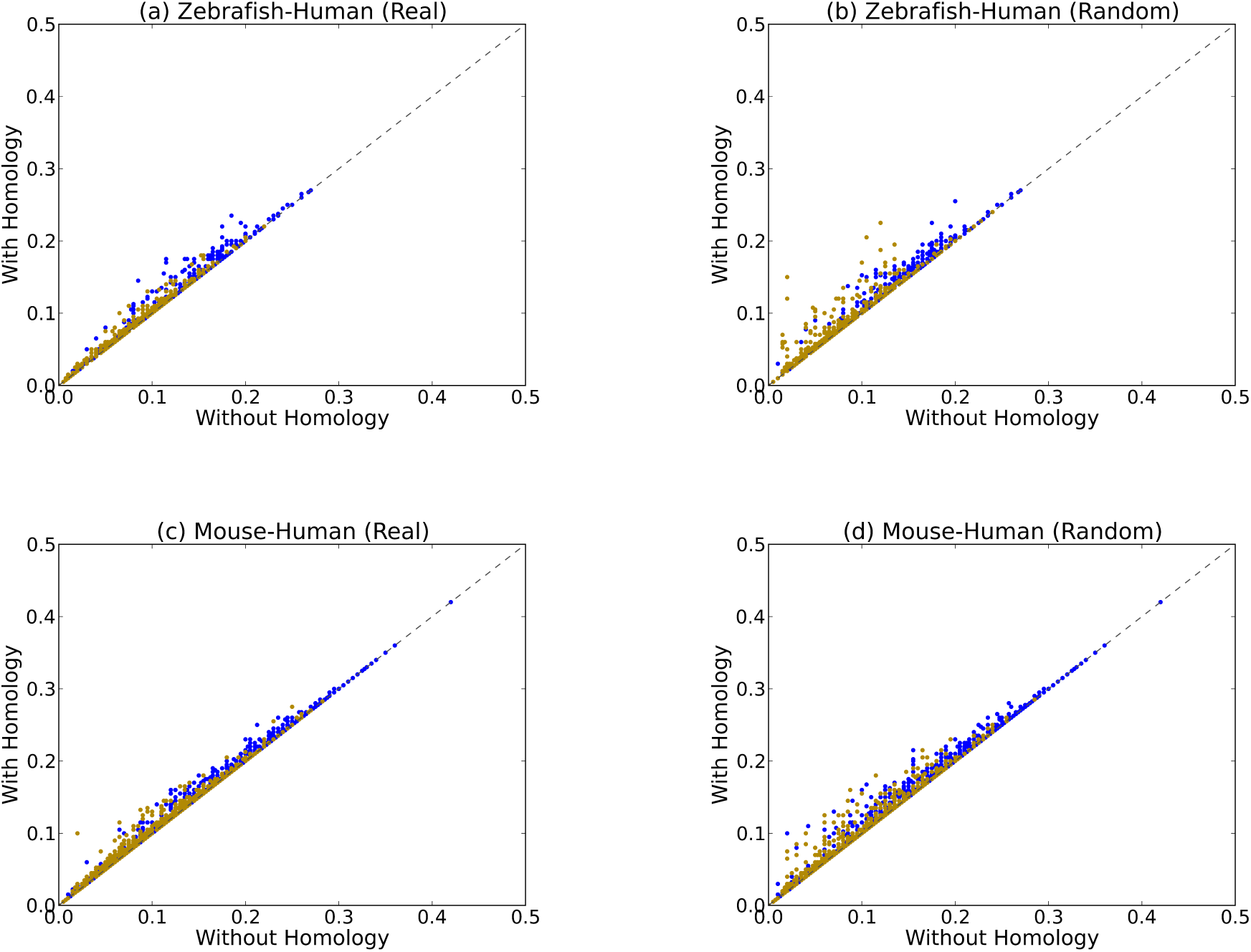
*S*_*IC,BP*_ for orthologs (blue) and non-orthologs (yellow) for the zebrafish-human (a,b) and mouse-human (c,d) species comparisons. The *x*-axis shows the scores without homology axioms (*R*) and the *y*-axis shows the scores for real (*H*) homology axioms (a and c) or random axioms (b and d).

**Table 4.** Differences in similarity between orthologs and non-orthologs upon adding either real or random homology axioms to *R*. *t*: one-tailed, unpaired *t*-statistic; df: degrees of freedom; ns: not significant; *δ_O_*, *δ_NO_*: mean percent increase ± 2 standard errors relative to *R* for orthologs and non-orthologs, respectively.

| species pair | | real homology ($H$) | random homology ($H'$) |
| --- | --- | --- | --- |
| zebrafish-human | $t$, df=1278 | $t = 2.36, p = 0.009$ | $t = 1.46, \text{ns}$ |
| | $\delta_O$ | $5.81 \pm 0.70$ | $4.85 \pm 0.98$ |
| | $\delta_{NO}$ | $4.88 \pm 0.72$ | $7.99 \pm 3.11$ |
| mouse-human | $t$, df=4066 | $t = -3.17, \text{ns}$ | $t = -2.16, \text{ns}$ |
| | $\delta_O$ | $2.44 \pm 0.27$ | $3.22 \pm 0.61$ |
| | $\delta_{NO}$ | $4.39 \pm 0.53$ | $5.83 \pm 0.81$ |

## 4. Discussion

We wished to measure the extent to which addition of homology axioms to an anatomy ontology affects the semantic similarity of phenotypes between distantly related species. The pattern whereby orthologs between distantly related species (zebrafish and human) show a greater increase in similarity than non-orthologs when real homology axioms are added provides evidence that the inclusion of homology improves biological accuracy. However, there are three caveats. One, the relative difference in score between orthologs and non-orthologs, while significant, is less then < 1%. Two, there was, unexpectedly, a larger increase in the similarity score for non-orthologs in the mouse-human comparison using real homology axioms. Third, non-orthologs had a greater increase in similarity for both species pairs when random homology axioms were added.

These results may be due to a combination of the hypothesized biological trend and a countervailing methodological artifact. First, the significant result for zebrafish-human with real homology axioms is consistent with the idea that the strength of the effect of including real homology axioms is in proportion to the evolutionary distance between the species pair. Second, the greater response of non-orthologs than orthologs in the three other comparisons, may stem from both real and faux homology axioms having a greater effect on semantic similarity when phenotypes are dissimilar, as can be seen in Figure 4. When the species are closely related and orthologs are already highly similar, or when the axioms are random, then non-orthologs, which are less similar to begin with, preferentially experience the gain in similarity.

Despite the noisiness of the trends overall, we can see examples of individual gene pairs for which homology axioms have a large effect that makes biological sense. One such pair is the human gene TFAP2A (NCBI:gene:7020), which is annotated to “Fusion of middle ear ossicles”, and the zebrafish gene *tfap2a* (ZFIN:ZDB-GENE-011212-6), annotated to “abnormal(ly) decreased length quadrate”. The homology between the quadrate, part of the jawbone of basal vertebrates, and the incus, a middle ear ossicle in mammals, is a textbook example of vertebrate evolution.^25^ When homology was excluded, “abnormal(ly) decreased length quadrate” was matched to “Micrognathia” and grouped under the relatively uninformative grouping class “*Q* and ‘inheres in’ some *bone of jaw*” with an *Sim_IC_* score of 0.32. When homology assertions were included, these annotations were subsumed under the grouping class “*Q* and ‘inheres in’ some (‘homologous to’ some *auditory ossicle*)” with an *Sim_IC_* score of 0.56.

Despite examples such as this, the modest effect of homology overall was unexpected. One explanation could be that so much anatomical homology is already implicit within the Uberon ontology that homology axioms are only needed in rare cases. In practice, it is difficult to extricate groupings in the ontology that are based on characteristics such as morphology, function, and shared development from those based on homology, potentially rendering some homology axioms redundant. Another explanation for the modest effect of homology is the relatively low number of homology assertions added to *H* that are not self-homologies, and the fact that only a subset of those assertions are relevant to the taxonomic groups compared here. It is not clear to what extent the results might be affected by homologies known in the literature that have yet to be curated by Bgee.

Our analysis focused on humans and two vertebrate model organisms for which abundant mutant phenotype data and a convenient set of anatomical homology statements are available. Given that the effect of homology seemed to be more pronounced in the zebrafish-human comparison than that of mouse, it would be of interest to examine species pairs with even more divergent body plans. Unfortunately, there are relatively few anatomical homology axioms linking vertebrates with model organisms outside the vertebrates, such as fruitflies and nematodes. Nonetheless, these results suggest that it would be worthwhile to explore the impact of “deeper” homology statements, either those sourced from the literature, or those derived computationally, such as by the phenolog approach.^26^ In future work, we intend to explore the impact of homology reasoning on measurement of semantic similarity for phenotypes that vary naturally among vertebrate lineages, such as those in the Phenoscape Knowledgebase.^27^ Independent of the use of homology axioms, some of the semantic similarity statistics that we examined showed relatively poor discrimination between orthologs and non-orthologs, suggesting the need to take a critical look at the biological accuracy of different phenotype semantic similarity measures.

